# First record of *Aedes albopictus* in the department of Magdalena, North of Colombia: implications for arboviral disease transmission

**DOI:** 10.1101/2025.11.19.689261

**Authors:** Erik Perdomo-Balaguera, Katiuska Ariza-Campo, Libardo Atencio, José A. Usme-Ciro, Gabriel Parra-Henao

## Abstract

*Aedes albopictus* is widely distributed throughout the world. It was introduced into the Americas in 1985 and is considered a vector for arboviruses such as dengue. The species has been reported in fifteen departments of the country.

With this brief communication, we aim to report the finding of *Ae. albopictus* in five municipalities in the department of Magdalena.

Between June and November 2025, adult mosquitoes were collected and breeding sites were inspected in the five municipalities. Both the collected adults and larvae were identified at the entomology unit of the Public Health Laboratory of the Department of Magdalena and confirmed by the entomology laboratory of the National Institute of Health. Larvae and adults of Ae. albopictus were found in all five municipalities.

The detection of *Ae. albopictus* in five municipalities in the department of Magdalena highlights the importance of strengthening continuous entomological surveillance strategies at the municipal and departmental levels in the country, especially in Magdalena and the surrounding municipalities.

## Introduction

*Aedes (Ae*.*) albopictus*, known as the Asian tiger mosquito, has spread from Asia to all continents except Antarctica, mainly during the last five decades. This species is predominant in several countries, mainly in Europe, where it has been considered the primary vector for transmission of some urban arboviruses, including *Orthoflavivirus dengue* (DENV), *Alphavirus chikungunya* (CHIKV), and *Orthoflavivirus zikaense* (ZIKV) (1,2). Due to its highly invasive nature in combination with global warming, this species has rapidly increased its distribution range, currently representing a serious concern for public health worldwide (3, 4).

This species has an ample adaptive capacity to natural and urban ecosystems due to its physiological characteristics such as diapause in eggs exposed to extreme temperatures and larvae that can occupy various types of natural breeding sites, like the axils of leaves, tree holes, bamboo, bromeliads, and artificial breeding sites like tires and water tanks (5).

The colonization of various countries around the world by *Ae. albopictus* is related to the passive transport of its eggs through the intense trade in tires and ornamental aquatic plants (6). Its introduction and spread in the Americas during the 1980s due to the rapid expansion of air and sea traffic, favorable environmental conditions for its reproduction, its adaptability to the same deposits that serve as breeding sites for *Ae. aegypti* in domestic and peridomestic environments, and poor entomological surveillance for early vector control (7,8).

In the Americas, this species has been reported in various countries in both the continent and Caribbean islands, from the USA (9) to Argentina (10). In Colombia, *Ae. albopictus* was first recorded in 1998 in an abundantly vegetated suburban area of the Leticia municipality, located in the Amazonas department (11). Since then, it has been recorded in 54 localities in 15 of the country’s departments, including the departments of Amazonas, Antioquia, Arauca, Caldas, Casanare, Cauca, Choco, Cundinamarca, Nariño, Putumayo, Quindio, Risaralda, Santander, Valle del Cauca (10), and more recently in the department of Cordoba (12), the last one corresponding to the first report for the Caribbean region of Colombia. In this brief communication, we report the presence of this species even further north in the country, in the department of Magdalena. Therefore, the distribution of the species covered by this report extends from the southernmost part of the country to the north.

## Methods

The collection of immature and adult stages of mosquitoes was performed as part of the routine entomological surveillance in the urban areas of the municipalities of Algarrobo (10°11′19″N, -74°03′36″), Guamal (9°08′41″N -74°13′48″), Nueva Granada (9°48′05.3″N - 74°23′37.8″), San Zenón ( 9°14′42″, -74°29′57″W)) and San Sebastián de Buenavista (9°14′21″, -74°21′07″), in the Magdalena department, located in North of Colombia.

In each municipality, house-to-house inspections were conducted in urban blocks selected by the local entomological surveillance teams, considering historical arboviral transmission and operational feasibility. All potential larval habitats were examined inside the house (intradomicile), around the house (peridomicile, within a radius of ∼10 m) and in adjacent areas (lots and surrounding spaces), including ground-level water-storage tanks, plastic drums, abandoned toilets, barrels and disposable containers such as tires, cans and bottles. Immature stages were collected using manual pipettes and dippers, placed in labelled vials and transported to the entomology laboratory of the Secretaría de Salud del Magdalena. Adult mosquitoes resting both indoors and outdoors were captured with entomological nets and mouth aspirators. Larvae and adults were examined under a stereomicroscope and identified using the morphological keys of Rueda (13) The main diagnostic characteristics for larvae and adults were identified. The diagnostic characters observed to identify the larvae were setae VII with a double or triple head, thorax with short lateral spines; teeth of the segment VIII comb without subapical spines (Figure 1). and, in adults, the clipping without spots of white scales, the thorax scutum with a narrow white median-longitudinal stripe, and median femur without longitudinal white stripe (figure 2) and identified as *Ae. albopictus*, This Identification was confirmed by the National Institute of Health of Colombia.

**Figure 1.**
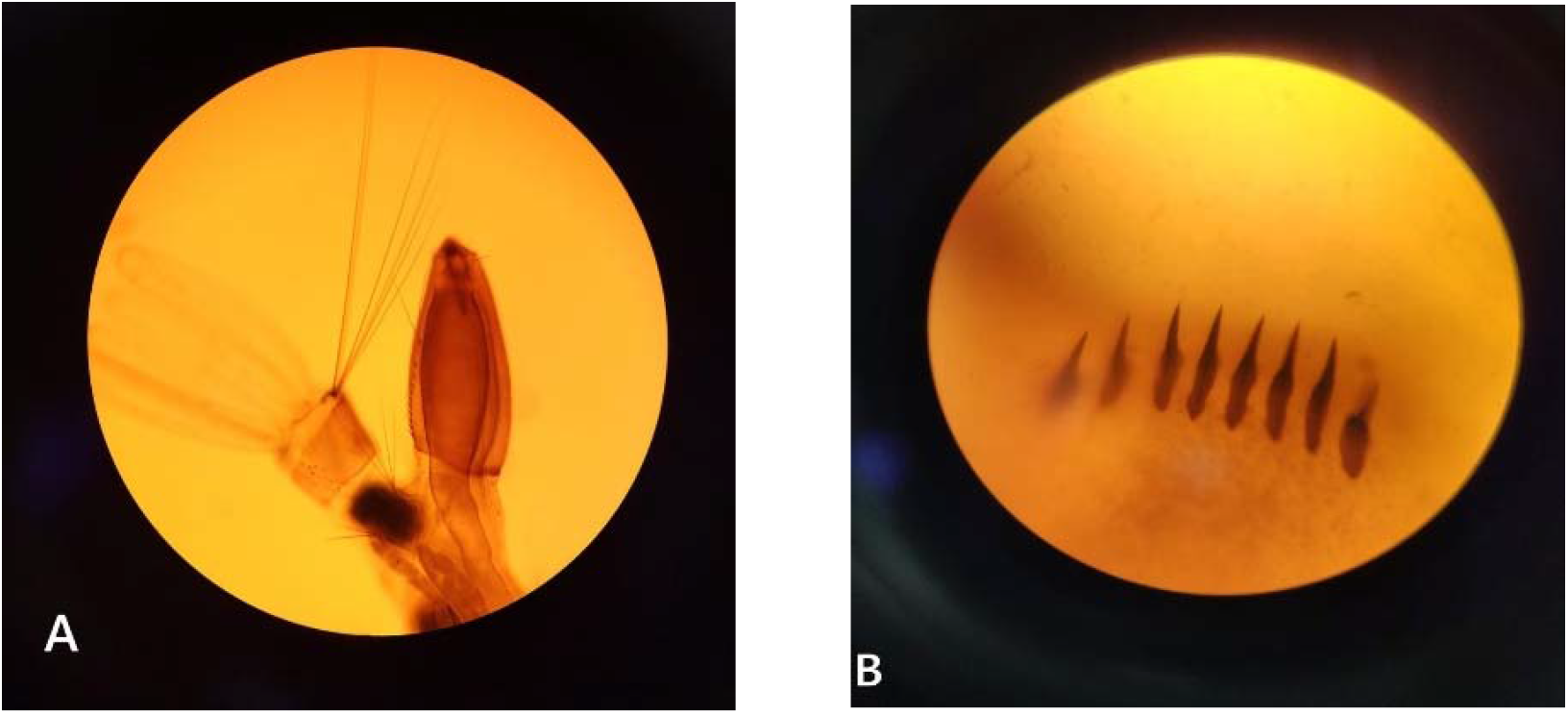
Diagnostic characteristics in Ae. albopictus larvae: A. Ventral brush (4-X) with four pairs of setae. B. Comb teeth straight spine-like without subapical spines.

**Figure 2.**
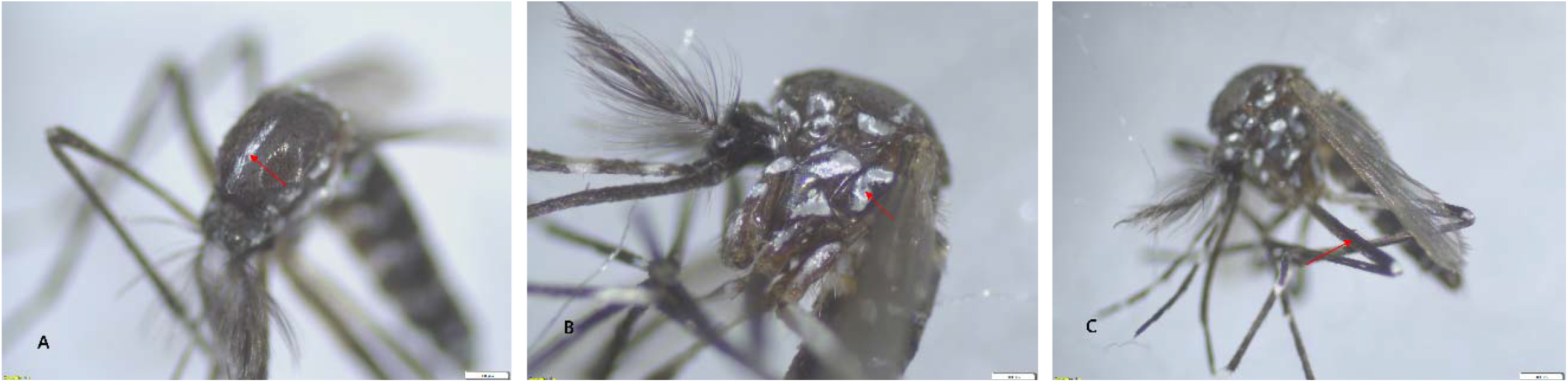
A) Thorax: Scutellum with a medial-longitudinal white stripe. B) Mesepimeron with non-separated white scales, forming a V-shaped white spot; C) Anterior portion of the medial leg femur without a longitudinal white stripe.

This finding means the geographical distribution of *Ae. albopictus* in Colombia is extending to 16 departments (figure 3) and this is the northernmost discovery in the country.

**Figure 3.**
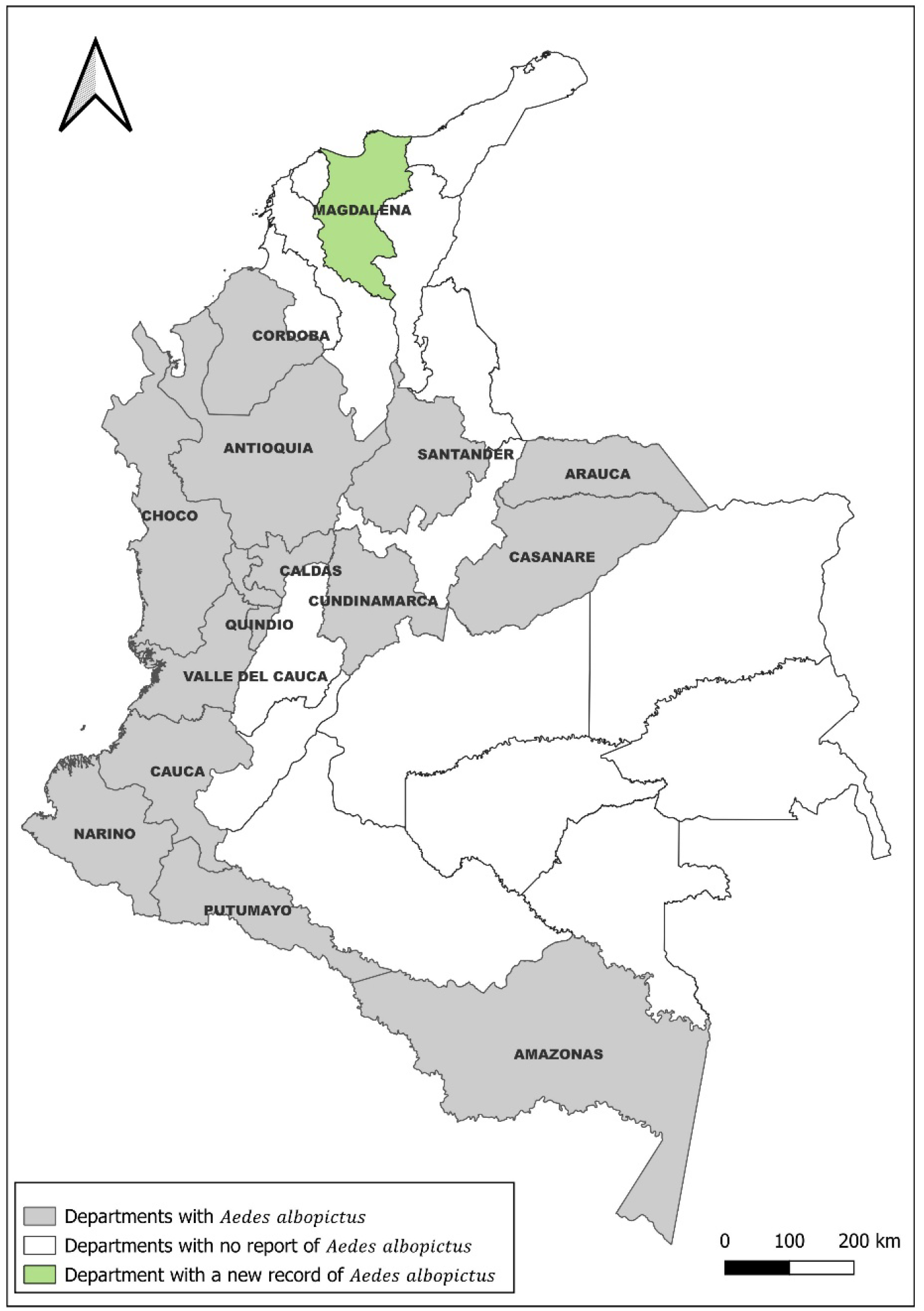
Geographical distribution of *Aedes albopictus* in Colombia. New record in Magdalena state in green.

In total, 10 adult and 64 larvae of *Ae. albopictus* were identified in the sampled municipalities; 16 individuals were collected in Algarrobo, 34 in Guamal, 6 in Nueva Granada, 10 in San Sebastián de Buenavista, and 8 in San Zenón (Table 1). The spatial distribution of the five municipalities where *Ae. albopictus* was detected within the department of Magdalena is shown in Figure 4. Immature stages of *Ae. albopictus* were detected exclusively in artificial containers associated with drinking water storage, solid waste, and larval traps. In Algarrobo, larvae were collected from discarded tires and various containers; in Guamal, from various low-lying tanks; in Nueva Granada, from larval traps installed in the town’s central park; in San Zenón, from tires and a ground-level water tank; and in San Sebastián de Buenavista, from various small containers and a ground-level tank. In several dwellings, these containers were uncovered, partially filled, and located in shaded areas with surrounding vegetation, providing suitable microhabitats with organic matter and relatively stable water levels.

**Table 1.** Municipal characteristics at collection sites and counts of identified *Aedes albopictus* males and females.

| Municipality | Month of collection | Urban area population | Altitude (m a.s.l.) | Climate |  | Adults |  | Larvae |
| --- | --- | --- | --- | --- | --- | --- | --- | --- |
|  |  |  |  | Temperature (°C) | Average monthly precipitation (mm) | ♂ | ♀ |  |
| Algarrobo | June | 14,294 | 24 | 27-28 | 150-200 | 6 | 4 | 6 |
| San Sebastián de Buenavista | October | 18,865 | 23 | >28 | 150-200 | — | — | 10 |
| San Zenón | October | 11,279 | 25 | 27-28 | 150-200 | — | — | 8 |
| Nueva Granada | November | 17,470 | 20 | >28 | 150-200 | — | — | 6 |
| Guamal | November | 25,312 | 24 | >28 | 200 | — | — | 34 |

**Figure 4.**
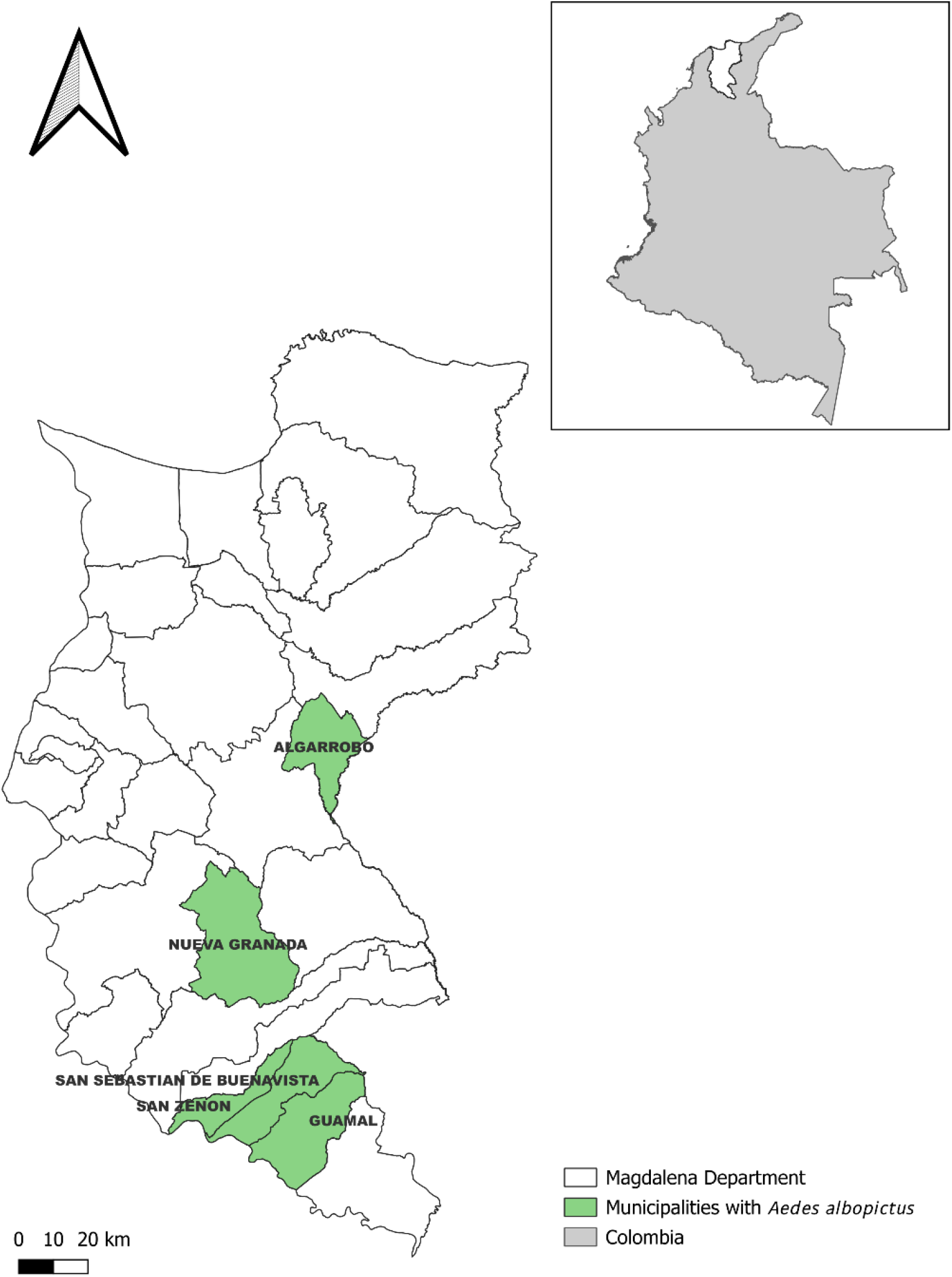
Municipalities of the department of Magdalena, northern Colombia, where Aedes albopictus was detected in 2025. Highlighted areas indicate the five municipalities with confirmed records (Algarrobo, San Sebastián de Buenavista, San Zenón, Nueva Granada and Guamal)

Since the first record of *Ae. albopictus* in the Amazonas department, in southern Colombia, and up to date, its geographical dispersion route has been indicated by reports in the country’s western, central and eastern departments, which means this is the second record for this species in north of Colombia, and the first report for the Magdalena department.

The accelerated distribution of *Ae. albopictus* in Colombia has been related to the availability of ecological habitats as breeding sites. Although the means that led to the colonization of *Ae. albopictus* in Magdalena department are unknown. The importance of human-mediated passive transport in the dispersal of the species is evident. In the Magdalena department, the introduction of *Ae. albopictus* appears to have occurred from the south, consistent with the species’ expansion pattern from the Amazon region toward northern Colombia; in this sense, it is consistent that Guamal, San Sebastián de Buenavista, San Zenón, Nueva Granada, and Algarrobo are the municipalities where its presence has been confirmed, arranged in a south–north belt along the Magdalena River and its road and fluvial connections, in line with a progressive northward spread mediated by passive transport (people, cargo, tires, and other containers). The municipalities of Algarrobo, San Sebastián de Buenavista, San Zenón, and Nueva Granada form a small river–road corridor that concentrates the movement of passengers, agricultural products, and solid waste, including used tires and other containers that retain water and can transport desiccation resistant eggs and immature stages. In this context, the road connections of Nueva Granada with these municipalities strengthen the network of exchange of goods and waste, integrating the municipality into the regional dynamics of vector dispersal. This network of connections, particularly the intense river traffic between Mompox and San Sebastián de Buenavista along the Magdalena River, provides repeated opportunities not only for the initial introduction but also for the reintroduction and local spread of *Ae. albopictus* within the department.

The rapid geographical expansion associated with this species’ biological and ecological characteristics, along with the persistent transmission of the DENV, the introduction of CHIKV in 2014 and ZIKV in 2015 to Colombia, and the circulation of other emergent and reemergent arboviruses in various ecological zones, demand a bigger effort in the entomological surveillance of this species, which is considered a potential vector of arboviral diseases of great importance in public health (12).

It is important to continue investigating the distribution of this vector in the department of Magdalena to better target and strengthen new entomological control strategies. Current surveillance and control activities in the department have focused almost exclusively on *Ae. aegypti*; however, our findings indicate that *Ae. albopictus* is now established in Magdalena and represents an additional public health threat, particularly given the presence of the Sierra Nevada de Santa Marta, an area with a history of yellow fever outbreaks. This finding confirms the establishment of the species in the department and underscores the need to adapt entomological surveillance and control strategies beyond *Ae. aegypti*, explicitly incorporating *Ae. albopictus* into routine risk assessment and intervention planning.

